# Disrupted maturation of white matter microstructure after concussion contributes to internalizing behavior problems in female children

**DOI:** 10.1101/2023.03.30.534745

**Authors:** Eman Nishat, Shannon E Scratch, Stephanie H Ameis, Anne L Wheeler

## Abstract

Some children that experience a concussion exhibit long-lasting emotional and behavioral problems post-injury, with greater rates of persistent problems in females. Establishing the contribution of (1) pre-existing behavioral problems and (2) disrupted maturation of the brain’s vulnerable white matter, to long-lasting behavioral problems has been a challenge due to a lack of pre-injury behavioral and imaging data. From the Adolescent Brain Cognitive Development Cohort, this study examined 204 11-12-year-old children who experienced a concussion after baseline data collection at age 9-10-years-old. Internalizing and externalizing behavioral problems were assessed with the Child Behavior Checklist. In 99 of these children with MRI data available, white matter microstructure was characterized in deep and superficial white matter by neurite density from restriction spectrum image modeling of diffusion MRI. Linear regressions modeled 1) post-concussion behavior symptoms controlling for pre-injury behavior, 2) the impact of concussion on white matter maturation, and 3) the contribution of deviations in white matter maturation to post-concussion behavior symptoms. When controlling for preinjury scores, post-injury internalizing and externalizing scores were higher in female but not male children with concussion compared to children with no concussion. Group comparisons of change in neurite density over two years reflecting white matter maturation demonstrated an age-dependent effect whereby younger female children had less change in neurite density over time than younger children with no concussion. In female children with concussion, less change in superficial white matter neurite density over time was associated with more internalizing behavior problems. These results suggest that in female children, concussions are associated with behavior problems beyond those that exist pre-injury, and injury to the brain’s vulnerable white matter may be a biological substrate underlying persistent internalizing behaviors.

## 2. Introduction

Concussions are an important public health concern with a high prevalence in children and adolescents (Dewan et al. 2016). Approximately 15-30% of children continue to experience symptoms more than one month after injury, with greater rates of persistent symptoms in females than males (Sheldrake et al. 2022; Zemek et al. 2016). Behavioral symptoms commonly observed following a concussion in childhood include externalizing behaviors (e.g., aggression, rule-breaking), and internalizing behaviors (i.e., anxiety, depression) (Ledoux et al. 2022; Li and Liu 2013). These symptoms can influence daily functioning and academic success and can lead to poor overall quality of life (Novak et al. 2016). However, the factors contributing to the onset and maintenance of these problems are unclear. Pre-existing behavioral problems are strong risk factors for persistent behavioral symptoms after concussion and may account for their apparent prevalence (Gornall et al. 2021; Zemek et al. 2016). At the biological level, developing white matter in the brain is highly susceptible to the stretching and shearing forces during concussions and its disruption and the resulting impact on brain function may contribute to behavioral symptoms.

Pre-existing behavioral problems complicate the assessment of post-concussion behavior symptoms in two ways. First, prior research has established pre-injury behavioral problems as a risk factor for experiencing a concussion in the first place (Goreth and Palokas 2019; Gornall et al. 2021), so cross-sectional comparisons with children who have not experienced a concussion are most often not matched on behavioral problems. Second, though pre-existing behavioral problems are significant predictors of long-lasting symptoms after concussions (Gagner et al. 2020; Max et al. 2015; Martin et al. 2020), the field has largely used parent-reported retrospective estimates which may not be reliable. Despite these caveats, studies looking at sexspecific effects show that female children with concussion are at greater risk of developing new behavioral problems post-concussion (Ellis et al. 2015), and are more likely to exhibit internalizing behavioral problems (Gornall et al. 2020). Overall, studies have not been able to draw definitive conclusions about the emergence of new behavioral problems after concussion due to the use of cross-sectional comparisons and absence of longitudinal data (Ledoux et al. 2022).

White matter in the brain is particularly vulnerable to primary and secondary injury caused by a concussion (Armstrong et al. 2016; Hulkower et al. 2013), which may disrupt its maturation that continues throughout childhood (Lebel, Treit, and Beaulieu 2019). Advanced diffusion MRI techniques, such as restriction spectrum imaging, can be used to detect microstructural properties of white matter that change over maturation, such as increased axon diameter, myelin content, and density of axons in tracts (White et al. 2013; Palmer et al. 2022). In a previous study, our research group found that female children who experienced a concussion before the age of 9 years old showed increased neurite density compared to children with no history of concussion, suggesting concussion is associated with disrupted white matter maturation (Nishat et al., 2022). However, this study, along with most pediatric concussion studies, is post-injury and cross-sectional, which limits the ability to directly assess changes in the developmental trajectory of white matter maturation. To investigate the direct effects of concussion on the developmental trajectory of white matter maturation, it is necessary to examine measures of white matter microstructure from both before and after the concussion. Using this information, we can more closely examine how disruption to white matter maturation contributes to behavioral symptoms after concussion.

This study leverages behavioral and imaging data acquired before the concussion in children to assess whether disruption of white matter maturation contributes to new behavior problems after a concussion. The sample consists of children from the Adolescent Brain Cognitive Development (ABCD) Study with pre-concussion data collected at age 9-10-years-old and post-concussion data collected at age 11-12-years-old who are compared to children from the ABCD study that did not experience a concussion. We first investigate whether post-injury internalizing and externalizing behavior problem scores are higher in children with concussion after accounting for preinjury problems. We next assess the impact of concussion on white matter maturation by examining change in neurite density derived from restriction spectrum image modeling of diffusion MRI over two years in deep and superficial white matter. Upon establishing the presence of behavioral problems that are a result of the concussion, we assess the contribution of disrupted white matter maturation to new behavioral problems. As our previous study in a distinct subsample of the ABCD Study indicated specific white matter impairment in females, all analyses are examined in a sex-stratified manner with the hypothesis that, in females, new behavioral problems would be associated with white matter disruption.

## 3. Methods

### 3.1. Participants and demographic data

This study examined a subset of children in the Adolescent Brain Cognitive Development (ABCD) Study, a large open-access developmental dataset of children aged 9-to-10-years-old at study entry (baseline) and 11-to-12-years-old at the two-year follow-up. Participants were recruited from 21 different sites across the United States of America (USA) and closely match the demographic characteristics of the USA population (Garavan et al. 2018) (https://abcdstudy.org/; https://abcdstudy.org/families/procedures/). Demographic data including age and sex assigned at birth were collected from the Physical Health measure completed by parents, and total combined family income and race and ethnicity were collected from the Parent Demographics Survey. History of medication taken in the two weeks before assessments were collected from the Parent Medications Survey Inventory Modified from PhenX. Any missing demographic data were imputed using the R Multivariate Imputation by Chained Equations (mice) package to avoid deletion of participants (Buuren and Groothuis-Oudshoorn 2011; R Development Core Team 2019). Siblings were randomly excluded from the sample to only include one sibling to remove any risk of Type 1 errors due to within-family nesting effects (Saragosa-Harris et al. 2021). Children that experienced a concussion after study entry and before the two-year follow-up were compared to children with no history of brain injury. Children with concussion were excluded from the sample if their injury occurred less than one month before the follow-up scan date to investigate changes in the post-acute phase after injury (n = 2). Signed informed consent was provided by all parents/or guardians and assent was obtained from children before participation in the study. The Research Ethics Board of The Hospital for Sick Children approved the use of ABCD data for this study.

### 3.2. Concussion

Details of the injury and concussion history were collected from the Modified Ohio State University Traumatic Brain Injury Screen – Short Version (Corrigan and Bogner 2007). This is a parent-report of children’s head injuries experienced since the last interview with questions about the mechanism of injury, the age at injury, and whether the child experienced any loss of consciousness or a gap in their memory. Children with concussion were identified according to the Summary Scores of Traumatic Brain Injury (TBI) derived from the screen and provided by the ABCD Study. Those that received a severity grading of ‘possible mild TBI’ (TBI without loss of consciousness but memory loss) or ‘mild TBI’ (TBI with loss of consciousness less than 30 minutes), were included in this study.

### 3.3. Behavioral measure

Measures of children’s emotional and behavioral problems were collected from the Child Behavior Checklist (CBCL). The CBCL was completed annually by parents and contains 118 questions rated on a 3-point Likert scale (0 = not at all true; 1 = somewhat true; 2 = very true) about the child’s behavior over the last six months. The questions are organized into two broad groupings of syndromes: internalizing behavior (sum of anxious/depressed, withdrawn/depressed, and somatic complaints scales) and externalizing behavior (sum of rule-breaking and aggressive behavior scales) (Achenbach and Ruffle 2000). This study examined children that had complete pre-injury baseline and post-injury follow-up behavioral information (concussion: n = 204, comparison group: n = 4981).

### 3.4. Magnetic resonance imaging (MRI)

#### 3.4.1. Acquisition and processing

MRI, including T1-weighted and diffusion-weighted imaging, was collected across 21 sites, on 28 different scanners, including Siemens Prisma, Philips, and GE 3T. MRI acquisitions, processing pipelines and quality control procedures are described in detail elsewhere (Hagler et al. 2019; Casey et al. 2018), and relevant details are summarized here. ABCD provided pre-processed region-based measures from multi-shell diffusion-weighted images acquired using 1.7 isotropic resolution, multiband EPI, slice acceleration factor 3, seven b=0 frames, and four b-values at 96 diffusion directions (six directions at b=500 s/mm2, 15 directions at b=1000 s/mm2, 15 directions at b=2000 s/mm2, and 60 directions at b=3000 s/mm2), compiled across subjects and summarized in a tabulated form. Diffusion MRI images were corrected for eddy current distortion, head motion, spatial, and intensity distortion, and registered to T1-weighted structural images using mutual information after coarse prealignment via within-modality registration to atlas brains (Wells et al. 1996; Hagler et al. 2019b). Surface-based nonlinear registration to the Desikan-Killiany atlas based on cortical folding patterns allowed for identification of superficial white matter beneath the cortical regions, and deep white matter tract segmentation and labeling was performed using AtlasTrack, a probabilistic atlas-based method (Hagler et al. 2019a).

#### 3.4.2. Quality control and exclusion criteria

All images underwent ABCD quality control procedures, including both automated checks for the completeness of the imaging series and to confirm that the number of files matched the expected number for each series on each scanner, and manual checks to identify poor image qualities or errors from motion correction. Participants were excluded if they had missing or failed quality control as rated by the ABCD Study team (concussion: n = 51, comparison group: n = 1106), missing imaging data (concussion: n = 33, comparison group: n = 773). Participants were also excluded if they had a diagnosis of epilepsy (concussion: n = 4, comparison group: n = 58), lead poisoning (concussion: n = 0, comparison group: n = 9), multiple sclerosis (n = 0), or cerebral palsy (concussion: n = 0, comparison group: n = 1), since the presence of these brain disorders may influence brain imaging. Children with concussion were excluded from the sample if their injury occurred before the baseline scan date (n = 17). Following these exclusions, the imaging analyses in this study examined participants with complete baseline and follow-up imaging data (concussion: n = 99, comparison group: n = 3024).

#### 3.4.3. Restriction spectrum imaging (RSI)

RSI models the relative proportion of restricted diffusion within intracellular space and hindered diffusion within extracellular space, further delineating restricted isotropic diffusion from restricted anisotropic diffusion. Developmental factors that contribute to the greater proportion of restricted isotropic diffusion in white matter include myelination and increases in axon diameter and axon density (Palmer et al. 2022). The RSI model was fit using a linear estimation approach as described by Hagler et al (2019). We used mean neurite density values derived from intracellular restricted diffusion and calculated for two categories of tissue: i) deep white matter, from 37 long-range major white matter tracts labeled using AtlasTrack (Hagler et al. 2009), and ii) superficial white matter, sub-adjacent to 68 regions of the cortex labeled using anatomical parcellation from the Desikan-Killiany atlas (Desikan et al. 2006).

To estimate and remove scanner effects from MRI measures, we used ComBat harmonization for longitudinal data (https://github.com/jcbeer/longCombat), with sex, age, and sociodemographic variables as covariates in the design matrix (Beer et al. 2020).

### 3.5. Statistical analyses

All analyses were conducted in R using the *stats* package for all linear regression models and *lme4* package for all linear mixed effect models. For each analysis, interaction effects were included to evaluate if concussion-specific effects were dependent on age or sex. This included a three-way group-by-sex-by-age interaction model, a two-way group-by-sex interaction model, a two-way group-by-age interaction model, and a model that included group, sex, and age as main effects. The Akaike Information Criterion (AIC) was used to identify the best-fit model. First, we report the best-fit model for each analysis. Next, we report sex-stratified analyses to investigate any sex-specific differences in females and males separately. All models controlled for sociodemographic variables including race/and ethnicity, combined family income, and puberty.

### 3.6. Experimental design

#### 3.6.1. Group comparisons of behavior

Linear regression models were generated to assess group differences in externalizing and internalizing behavior scores at follow-up between children with and without concussion. These models included baseline internalizing and externalizing behavior scores to control for any pre-injury differences.

#### 3.6.2. Group comparisons of change in neurite density over time

White matter maturation over the two years was investigated by calculating the change in white matter neurite density in deep white matter and superficial white matter from baseline to follow-up normalized by time since baseline. Linear mixed effect models were generated to assess group differences in the change in neurite density over time between children with and without concussion. These models included baseline neurite density as a fixed effect covariate and scanner as a random effect.

##### 3.6.2.1. Exploratory post-hoc analyses

###### 3.6.2.1.1. Group comparisons of change in white matter within age quartiles

To explore any age interactions driving group differences between children with and without concussion, we divided children based on age quartiles and compared neurite density change differences between children with and without concussion in the top and bottom age quartiles. Linear mixed-effect models were set up as described above.

###### 3.6.2.1.2. Association between change in white matter and injury variables in children with concussion

In children with concussion, we investigated the relationship between injury variables and change in neurite density in tissue categories where group differences in white matter maturation were detected. Linear mixed effect models included injury mechanism, number of injuries, number of injuries with loss of consciousness, number of injuries with post-traumatic amnesia, age at injury, time since first injury, and baseline neurite density as fixed effect covariates, and scanners as random effect.

#### 3.6.3. Association between change in white matter and behavior in children with concussion

In white matter tissue categories where significant differences in maturation were detected in children with concussion, we calculated a deviation score to characterize white matter maturation in the concussion group relative to the expected trajectory of white matter maturation as seen in the comparison group. First, mean and standard deviation of the comparison group’s neurite density change scores were calculated. Next, z-scores were computed for each participant within the concussion group based on the mean and standard deviation of the comparison group. These z-scores were defined as the deviation score. Positive deviation scores indicate more maturation in an individual in the concussion group and negative deviation scores indicate less maturation in an individual in the concussion group compared to the comparison group. Linear regression models were generated to investigate the relationship between deviation of white matter maturation and post-injury internalizing and externalizing behavior scores. These models included pre-injury internalizing and externalizing behavior scores as a fixed effect covariate.

### 3.7. Data availability

All data used in this study is from the ABCD Study’s Curated Annual Release 2.0 and 4.0 (https://data-archive.nimh.nih.gov/abcd).

### 3.8. Code accessibility

Code for implementing the analyses presented here is found at https://github.com/eman-nishat/eman-proj/tree/main/pre-postinjury-group-analyses. Requests to access the derived data in this study should be directed to Anne Wheeler.

## 4. Results

### 4.1. Participants

The demographics and characteristics of the concussion and comparison groups are presented in **Table 1** and the injury characteristics for the concussion group are presented in **Table 2**. Demographics and injury characteristics of the concussion group with MRI data available are presented in **Supplementary Table 1**. The concussion group contained a higher proportion of males, was slightly older and had higher family income on average compared to the comparison group. The concussion group also contained more non-hispanic white participants and less hispanic and non-hispanic black participants. These variables were all controlled for in the following statistical analyses. Most concussions resulted from a fall with one quarter of participants experiencing loss of consciousness and most experiencing memory loss.

**Table 1.** Demographics and characteristics of the concussion and comparison groups

|  | Comparison Group<br>(N=4981) | Concussion<br>(N=204) | p-value |
| --- | --- | --- | --- |
| Sex |  |  |  |
| Female | 2509 (50.4%) | 78 (38.2%) | <0.001 |
| Male | 2472 (49.6%) | 126 (61.8%) |  |
| Age at Baseline (in months) |  |  |  |
| Mean (SD) | 119 (7.35) | 120 (7.29) | 0.0128 |
| Median [Min, Max] | 119 [108, 131] | 121 [108, 131] |  |
| Age at Follow-Up (in months) |  |  |  |
| Mean (SD) | 143 (7.72) | 145 (7.58) | 0.00469 |
| Median [Min, Max] | 143 [127, 168] | 146 [129, 165] |  |
| Puberty at Follow-Up |  |  |  |
| Prepubescence | 2470 (49.6%) | 115 (56.4%) | 0.0675 |
| Pubescence | 2511 (50.4%) | 89 (43.6%) |  |
| Combined Family Income |  |  |  |
| <\$50K | 1533 (30.8%) | 31 (15.2%) | <0.001 |
| \$100K+ | 2012 (40.4%) | 103 (50.5%) | |
| \$50-99K | 1436 (28.8%) | 70 (34.3%) | |
| Race/Ethnicity |  |  |  |
| Asian | 131 (2.6%) | 5 (2.5%) | <0.001 |
| Hispanic | 1092 (21.9%) | 27 (13.2%) |  |
| Non-Hispanic Black | 684 (13.7%) | 13 (6.4%) |  |
| Non-Hispanic White | 2606 (52.3%) | 134 (65.7%) |  |
| Other/Multi-Racial | 468 (9.4%) | 25 (12.3%) |  |
| Pre-Injury Internalizing Behaviour Raw Score |  |  |  |
| Mean (SD) | 4.70 (5.18) | 6.11 (6.47) | 0.00243 |
| Median [Min, Max] | 3.00 [0, 51.0] | 4.50 [0, 36.0] |  |
| Pre-Injury Internalizing Behaviour T-Score |  |  |  |
| Mean (SD) | 47.8 (10.3) | 50.6 (11.0) | <0.001 |
| Median [Min, Max] | 48.0 [33.0, 93.0] | 50.0 [33.0, 84.0] |  |
| Pre-Injury Externalizing Behaviour Raw Score |  |  |  |
| Mean (SD) | 3.96 (5.41) | 5.01 (6.27) | 0.0187 |
| Median [Min, Max] | 2.00 [0, 48.0] | 2.50 [0, 29.0] |  |
| Pre-Injury Externalizing Behaviour T-Score |  |  |  |
| Mean (SD) | 44.8 (10.0) | 46.6 (10.9) | 0.0181 |
| Median [Min, Max] | 44.0 [33.0, 83.0] | 45.0 [33.0, 74.0] |  |
| Post-Injury Internalizing Behaviour Raw Score |  |  |  |
| Mean (SD) | 4.66 (5.36) | 6.32 (6.33) | <0.001 |
| Median [Min, Max] | 3.00 [0, 38.0] | 4.00 [0, 40.0] |  |
| Post-Injury Internalizing Behaviour T-Score |  |  |  |
| Mean (SD) | 47.2 (10.4) | 50.7 (10.5) | <0.001 |
| Median [Min, Max] | 46.0 [33.0, 82.0] | 50.0 [33.0, 81.0] |  |
| Post-Injury Externalizing Behaviour Raw Score |  |  |  |
| Mean (SD) | 3.60 (5.15) | 4.95 (6.27) | 0.00268 |
| Median [Min, Max] | 2.00 [0, 46.0] | 3.00 [0, 35.0] |  |
| Post-Injury Externalizing Behaviour T-Score |  |  |  |
| Mean (SD) | 43.9 (9.52) | 46.3 (10.3) | 0.0012 |
| Median [Min, Max] | 43.0 [33.0, 82.0] | 46.0 [33.0, 74.0] |  |

**Table 2.** Injury characteristics of the concussion group

|  | **F**<br>**(N=78)** | **M**<br>**(N=126)** | **Total**<br>**(N=204)** |
| --- | --- | --- | --- |
| **Age at Injury (in months)** |  |  |  |
| Mean (SD) | 127 (10.7) | 128 (15.1) | 128 (13.6) |
| Median [Min, Max] | 132 [108, 144] | 132 [12.0, 156] | 132 [12.0, 156] |
| **Mechanism of Injury** |  |  |  |
| ER | 15 (19.2%) | 23 (18.3%) | 38 (18.6%) |
| Fall | 50 (64.1%) | 82 (65.1%) | 132 (64.7%) |
| MVC | 9 (11.5%) | 9 (7.1%) | 18 (8.8%) |
| Other LOC | 4 (5.1%) | 6 (4.8%) | 10 (4.9%) |
| Blast | 0 (0%) | 2 (1.6%) | 2 (1.0%) |
| Fight | 0 (0%) | 3 (2.4%) | 3 (1.5%) |
| Repeated Head Impact | 0 (0%) | 1 (0.8%) | 1 (0.5%) |
| **Loss of Consciousness** |  |  |  |
| No | 59 (75.6%) | 93 (73.8%) | 152 (74.5%) |
| Yes | 19 (24.4%) | 33 (26.2%) | 52 (25.5%) |
| **Memory Loss** |  |  |  |
| No | 7 (9.0%) | 17 (13.5%) | 24 (11.8%) |
| Yes | 71 (91.0%) | 109 (86.5%) | 180 (88.2%) |
| **Time Since Injury (in months)** |  |  |  |
| Mean (SD) | 12.7 (14.9) | 11.2 (11.9) | 11.8 (13.1) |
| Median [Min, Max] | 10.0 [1.00, 136] | 10.0 [1.00, 130] | 10.0 [1.00, 136] |

\*ER = Injury Resulting in Emergency Room Visit, MVC = Motor Vehicle Crash, LOC = Loss of Consciousness

### 4.2. Group comparisons of behavior post-injury, controlling for preinjury scores

Children with concussion had higher internalizing and externalizing behavior scores both pre- and post-injury than children with no history of concussion (**Table 1**). To investigate whether the differences in scores post-injury are due to pre-existing behavioral problems, we first compared post-injury scores between groups, controlling for pre-injury scores.

#### 4.2.1. Internalizing behavior

The best-fit model carried 52% of the cumulative model weight and included a group-by-sex interaction. The group-by-sex interaction showed a trend level effect on internalizing behavior scores at follow-up when controlling for baseline scores (B = −1.099392, *p* = .07525).

Sex-stratified analyses showed female children with concussion had higher follow-up internalizing behavior scores compared to females with no concussion (B = 1.324046, *p =* .00862), and no differences in males (B = 0.29764, *p* = .4237) (**Figure 1A**).

**Figure 1A.**
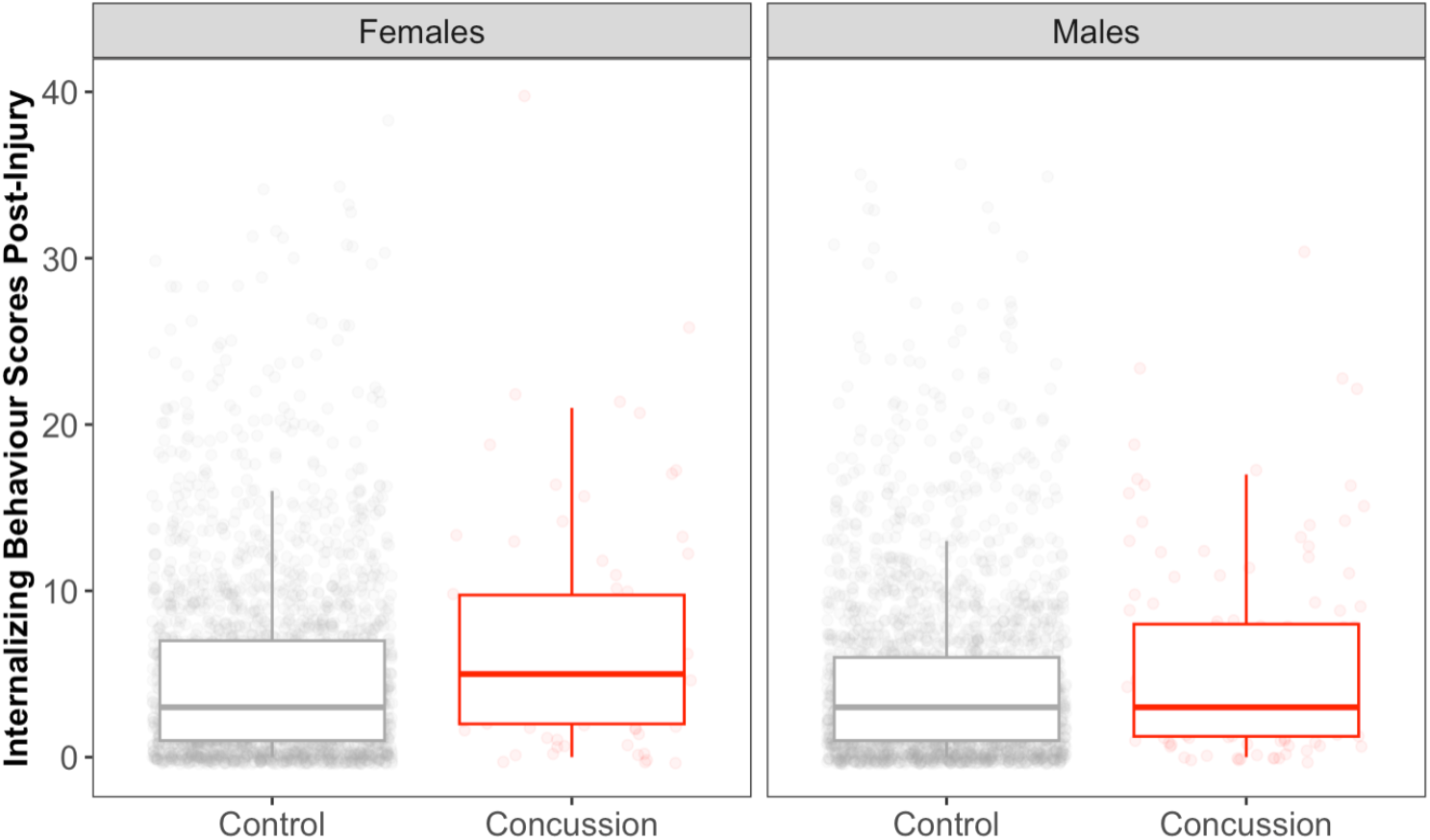
Group differences in internalizing behavior scores post-injury between concussion and comparison groups, and sex

#### 4.2.2. Externalizing behavior

The best-fit model carried 52% of the cumulative model weight and did not include any interaction terms. Main effect of group showed higher externalizing behavior scores at follow-up in children with concussion than children with no history of concussion (B = 0.651178, *p* = .016269).

Sex-stratified analyses showed higher scores in females with concussion than females with no history of concussion (B = 0.896602, *p* = .03078), and no differences in males (B = 0.44975, *p* = .21383) (**Figure 1B**).

**Figure 1B.**
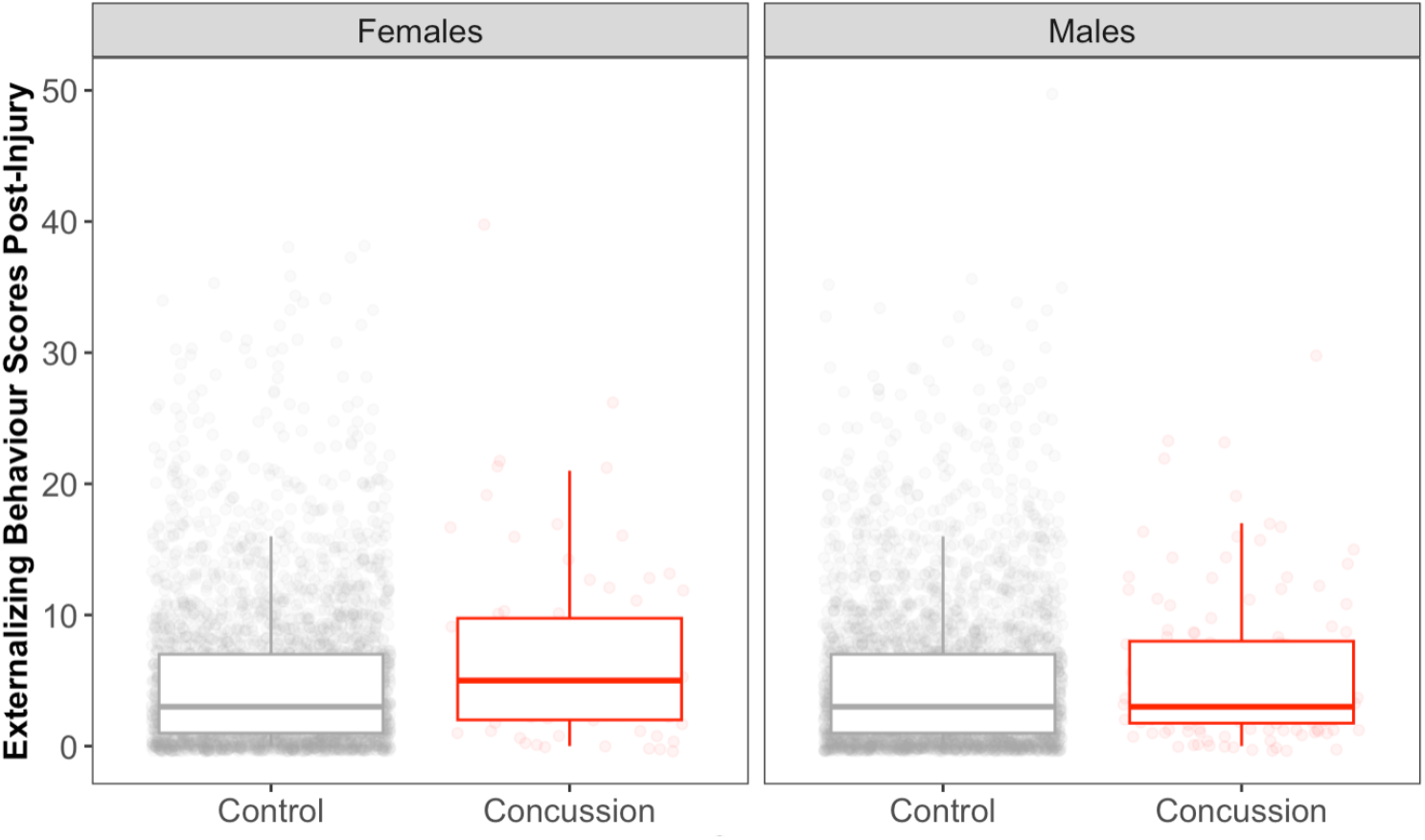
Group differences in externalizing behavior scores post-injury between concussion and comparison groups, and sex

### 4.3. Group comparisons of change in neurite density over time

To investigate whether concussion impacts white matter maturation over two years, we compared change in neurite density between groups, controlling for pre-injury neurite density (**Table 3**).

**Table 3.** Change in neurite density over two years in the concussion and control groups

| Change in Neurite Density Over Two Years | Comparison Group<br>(N=3024) | Concussion<br>(N=99) | p-value |
| --- | --- | --- | --- |
| Deep White Matter |  |  |  |
| Mean (SD) | 0.000398 (0.000895) | 0.000412 (0.000587) | 0.827 |
| Median [Min, Max] | 0.000391 [-0.00745, 0.0119] | 0.000374 [-0.00116, 0.00217] |  |
| Superficial White Matter |  |  |  |
| Mean (SD) | 0.000589 (0.00119) | 0.000547 (0.00119) | 0.73 |
| Median [Min, Max] | 0.000569 [-0.0125, 0.00821] | 0.000520 [-0.00608, 0.00479] |  |

#### 4.3.1. Deep White Matter

In both children with (**Figure 3A**) and without concussion (**Figure 3B**), there was an increase in neurite density of the deep white matter between the two scans over time (i.e., baseline to two-year follow-up) (**Table 3**).

**Figure 3.**
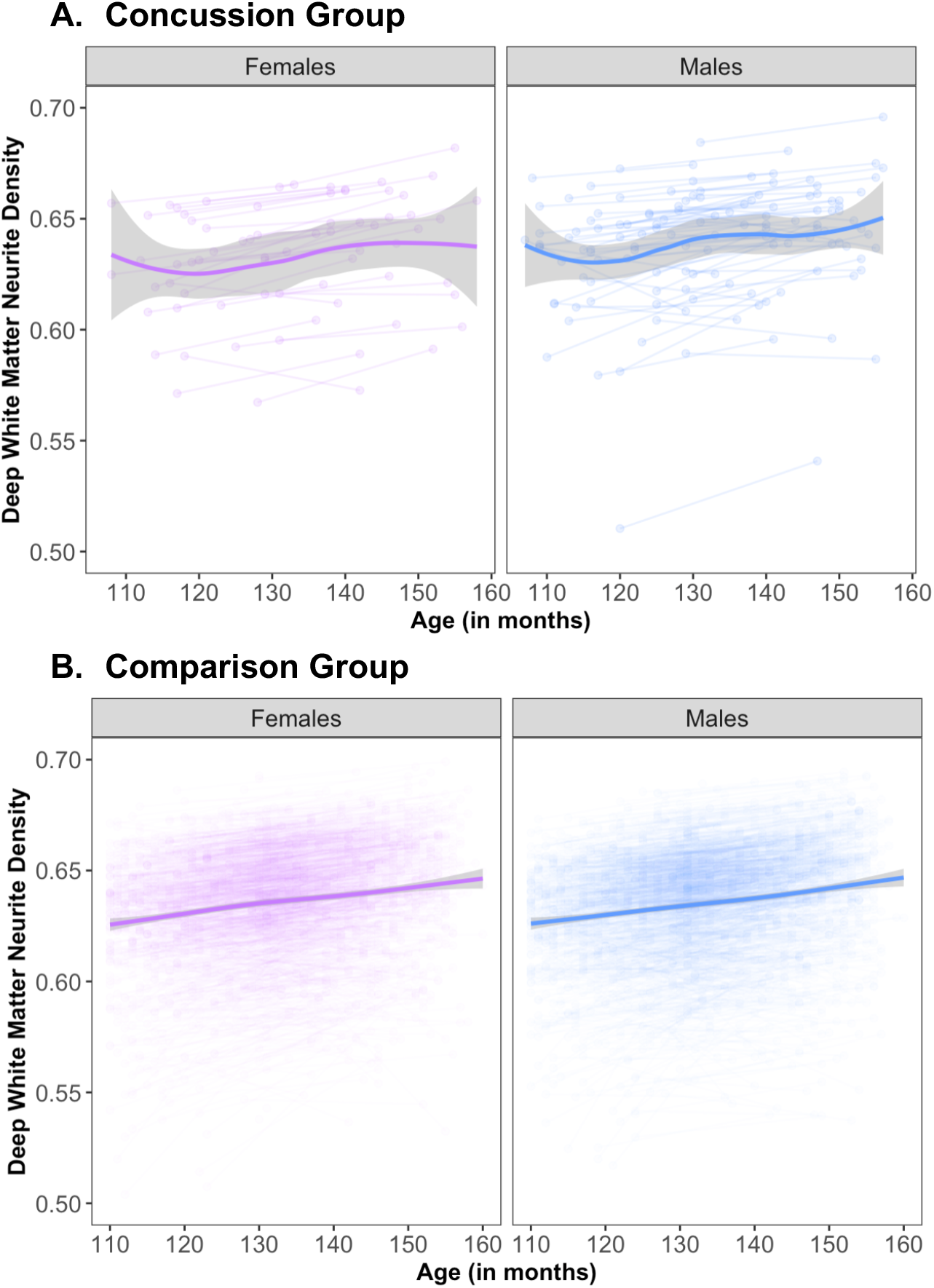
Neurite density at the baseline and followup scans reflecting deep white matter maturation over the two-year period in males and females (**A**) with and (**B**) without concussion.

When examining change in neurite density within deep white matter the best-fit model carried 56% of the cumulative model weight and did not include any interaction terms. Main effect of group did not show any significant differences in change in neurite density over time between children with and without a history of concussion (B = 1.019e-057.092e-06, *p* = .92727).

Sex-stratified analyses did not show any sex-specific group differences in females (B = 1.529e-05, *p* = .9021) or male (B = 1.043e-05, *p =* .91872) children with concussion.

#### 4.3.2. Superficial White Matter

In both children with (**Figure 4A**) and without concussion (**Figure 4B**), there was an increase in neurite density of the superficial white matter between the two scans (**Table 3**).

**Figure 4.**
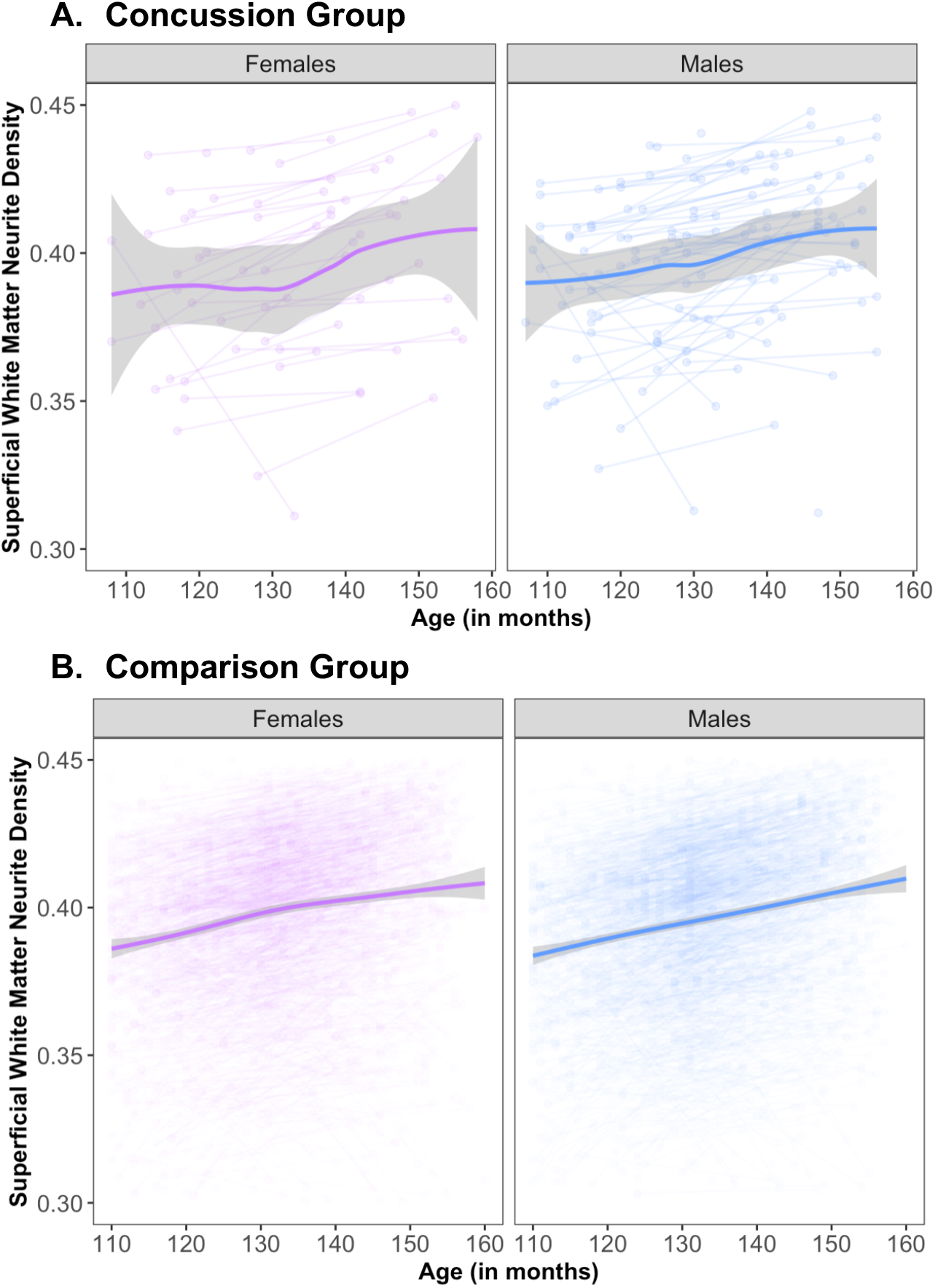
Neurite density at the baseline and followup scans reflecting superficial white matter maturation over the two-year period in males and females (**A**) with and (**B**) without concussion.

When examining change in neurite density, the best-fit model carried 50% of the cumulative model weight and included a group-by-age interaction (B = 3.270e-05, *p* = .02917). Both children with and without a history of concussion showed more change in superficial white matter neurite density with increasing age, with a more pronounced age-dependent effect in the concussion group (B = 3.926e-05, *p* = .01659) than in the comparison group (B = 7.746e-06, *p* = .00323) (**Figure 5)**.

**Figure 5.**
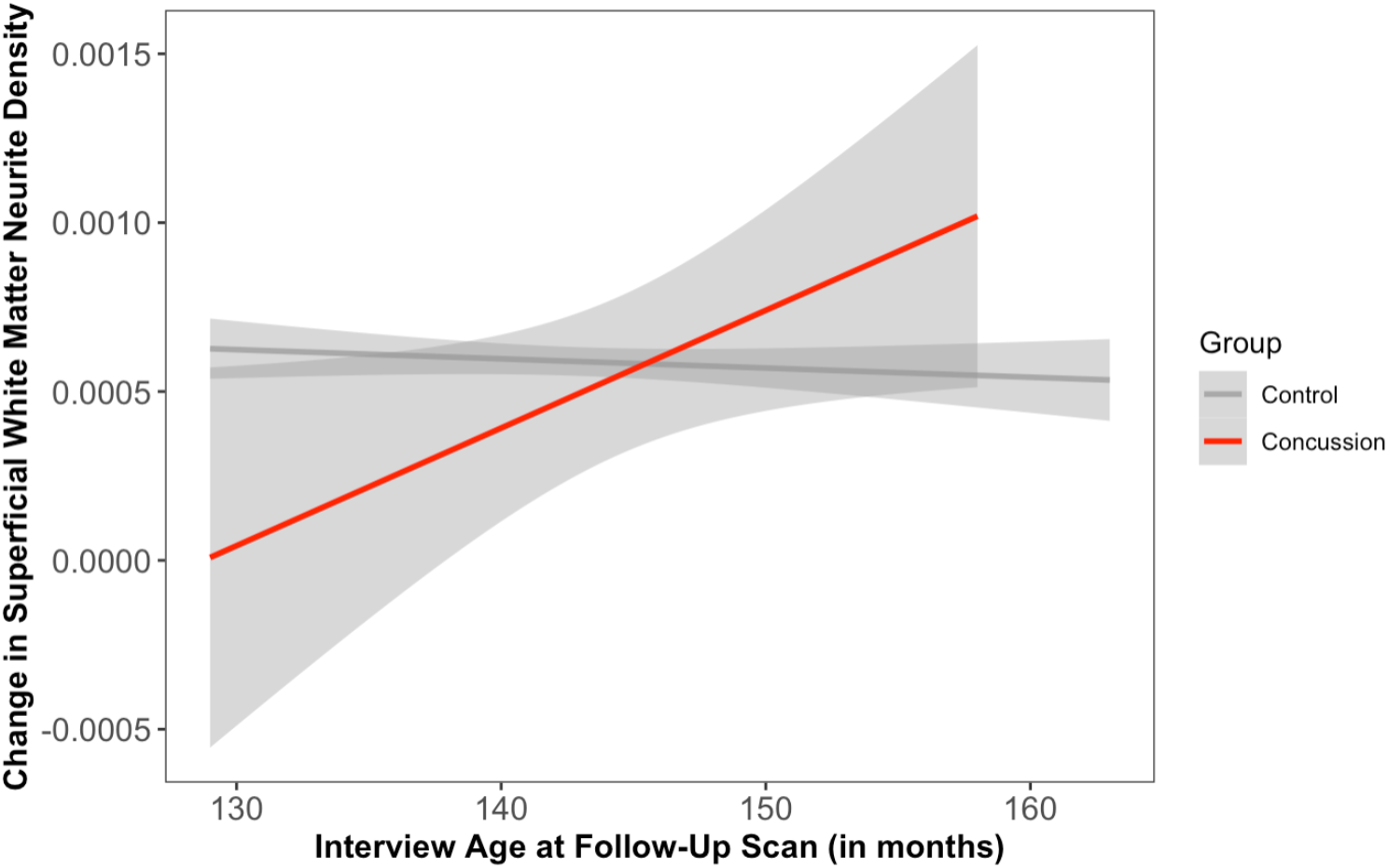
Regression lines showing group-by-age interaction when comparing change in superficial white matter neurite density between concussion and comparison groups.

Sex-stratified analyses with the group-by-age-interaction term showed that female children contributed most to this age dependent alteration in maturation (interaction B = 5.023e-05, *p* = .04779), but there was no significant interaction in males (B = −3.245e-03, *p* = .234354).

#### 4.3.3. Group comparisons of change in white matter within age quartiles

##### 4.3.3.1. Superficial White Matter

To further probe the group-by-age interaction observed when comparing change in superficial white matter neurite density between children with and without concussion, we compared neurite density change differences between children with and without concussion in the top and bottom age quartiles and found that the youngest children with concussion had less change in neurite density compared to children with no concussion (B = −7.764e-04, p = .0130) and there were no differences in the oldest children (B = −4.050e-05, p = .823).

Sex-specific differences were also reflected in the top and bottom age quartile group comparisons where younger females but not males with concussion showed less change in neurite density than children without concussion (F: B = −1.636e-03, p = .00166; M: B = −4.234e-04, p = .292). There were no significant differences in the oldest females or males with and without concussion (F: B = 2.011e-04, p = .542; M: B = −1.396e-04, p = .528).

#### 4.3.4. Association between change in neurite density and injury variables in children with concussion

To determine the relationship between superficial white matter maturation and injury severity, we investigated the influence of various injury factors on change in superficial white matter neurite density over time in the children with concussion.

##### 4.3.4.1. Superficial White Matter

There was a significant effect of time since injury, such that children with a more recent injury had greater change in superficial white matter neurite density over time (B = −6.821e-05, *p* = .02939) (**Table 4**).

**Table 4.** Association between change in neurite density and injury variables in children with concussion

|  | Group |  | Female |  | Male |  |
| --- | --- | --- | --- | --- | --- | --- |
|  | *Estimates* | *p-value* | *Estimates* | *p-value* | *Estimates* | *p-value* |
| Age at Injury (in months) | -0.00 | 0.149 | -0.00 | 0.355 | -0.00 | 0.265 |
| Mechanism of Injury - Fall | 0.00 | 0.706 | -0.00 | **0.025** | 0.00 | 0.149 |
| Mechanism of Injury - Fight | -0.00 | 0.751 |  |  | -0.00 | 0.854 |
| Mechanism of Injury - MVC | 0.00 | 0.101 | -0.00 | 0.618 | 0.00 | 0.064 |
| Mechanism of Injury - Other LOC | 0.00 | 0.916 | -0.00 | 0.222 | 0.00 | 0.523 |
| Mechanism of Injury - Repeated Head Impact | 0.00 | 0.415 |  |  | 0.00 | 0.382 |
| Time Since Injury | -0.00 | **0.029** | -0.00 | 0.659 | -0.00 | 0.058 |
| Loss of Consciousness (LOC) | 0.00 | 0.808 | 0.00 | 0.501 | -0.00 | 0.585 |
| Memory Loss | -0.00 | 0.857 | -0.00 | 0.266 | -0.00 | 0.646 |

Sex-stratified analyses showed injury from falls had a significant effect, such that female children that experienced a concussion from a fall had less change in superficial white matter neurite density over time (B = −9.139e-04, *p =* .04456) (**Table 4**)..

### 4.4. Association between deviation from change in superficial white matter neurite density and behavior scores in children with concussion

Female children with concussion had higher internalizing and externalizing behavior scores post-injury than children with no history of concussion, when controlling for pre-injury scores. This suggests that a disruption to underlying brain maturation may influence new behavioral problems in females. Indeed, age dependent alterations in superficial white matter maturation were detected in females as well. To examine how deviations from the expected trajectory of superficial white matter maturation in children with concussion may contribute to behavioral problems, we investigated the association between deviation from normative change in neurite density and behavior scores.

#### 4.4.1. Internalizing behavior

The best-fit model carried 79% of the cumulative model weight and included a deviation-score-by-sex interaction (B = 2.15242, *p* = .01757).

Sex-stratified analyses showed a significant relationship between deviation in superficial white matter neurite density change over time and post-injury internalizing behavior scores in females (B = −1.83158, *p* = .02839). Less maturation in superficial white matter neurite density in female children that had a concussion was associated with higher internalizing behavior scores after injury. This effect was consistent even when we removed four participants with extreme values, defined as participants with z-scores less than −2.5 (n = 2) or more than 2.5 (n = 2) (B = −4.21640, *p* = .04037). There were no significant relationships in males (B = −0.17759, *p* = .80244) (**Figure 6**).

**Figure 6.**
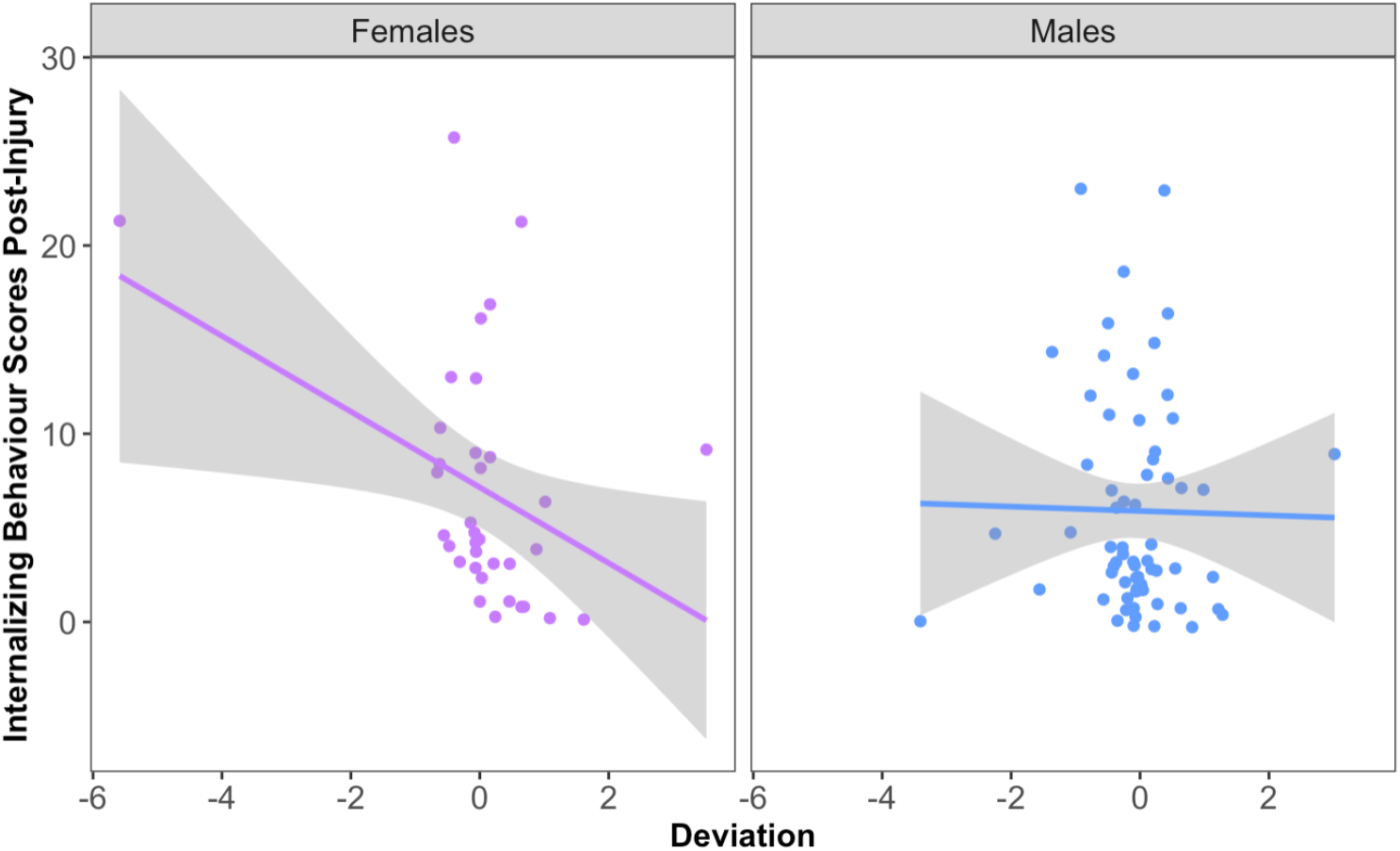
Relationship between deviation from change in superficial white matter neurite density and internalizing behavior problems

#### 4.4.2. Externalizing behavior

The best-fit model carried 62% of the cumulative model weight and did not include any significant interaction terms. There was no significant relationship between deviation from change in superficial white matter neurite density over time and follow-up externalizing behavior scores (B = −0.51800, *p* = .280197).

Sex-stratified analyses did not show any significant relationship between deviation from change in superficial white matter neurite density over time and follow-up externalizing behavior scores in females (B = −0.63923, *p* = .2902) or male (B = 0.38792, *p* = .442070) children with concussion.

## 5. Discussion

This present study investigated the contribution of pre-existing behavioral problems and disruption of white matter maturation to persistent behavioral problems after concussion in children aged 11-12-years-old in the ABCD Study. The findings confirm that children with concussion have more internalizing and externalizing behavior problems both before and after injury, compared to children without concussion. Interestingly, when examining the group differences in post-injury behavioral problems while controlling for pre-injury behavioral problems, female children had worse internalizing and externalizing behavior scores compared to females without concussion, while there were no differences in males. This supports previous literature suggesting that concussion may have negative effects on behavioral outcomes in children (Gornall et al. 2021; Li and Liu 2013; Schachar, Park, and Dennis 2015). However, consistent with previous research suggesting that females may be more susceptible to the negative effects of concussion on behavior (Gornall et al. 2020), our findings in the ABCD sample suggest that postconcussion behavior problems in males may not be due to the concussion and instead reflect pre-existing behavioral problems.

Next, the impact of concussion on white matter maturation was assessed by computing changes in neurite density over the two years between ages 9-10-years-old and 11-12-years-old. Neurite density from RSI is a specific measure of cylindrically restricted intracellular diffusion which is more sensitive to maturational changes than measured from fractional anisotropy derived from the more conventional diffusion tensor model (Zhao et al. 2021; Genc et al. 2017). Neurite density generally increased over time, consistent with increased myelination, axon diameter, and axon packing density which occurs over this developmental period (Palmer et al. 2022). When white matter change in the children with concussion was compared to children without concussion, there was a significant age-dependent group effect in superficial white matter. This age-dependent group effect appeared to be driven by reduced maturation in younger female children. Superficial white matter consists of short-range association fibers that connect adjacent gyri directly beneath the cortical surface (Wu et al. 2013). These fibers are late to myelinate in development and may be more impacted by concussion (Barkovich 2000). An earlier study by our research group showed that female children that experienced a concussion before the age of 9-10-years-old had higher neurite density indicating potentially enhanced maturation of superficial white matter. This earlier study examined a distinct subset of children from the ABCD Cohort that experienced a concussion before the baseline assessment so preinjury variables were not available and the comparisons were cross-sectional (Nishat et al., 2022). Taken together, these findings suggest that the effects of concussion on white matter maturation in females may depend on the age of injury, with different effects observed depending on the developmental stage of the brain. Furthermore, injury early in life may alter the maturational trajectory of white matter, whereby premature maturation is followed by slowed maturation, though future longitudinal studies would need to evaluate this more robustly.

Interestingly, when investigating the relationship between deviation from typical white matter maturation and behavioral problems in children with concussion, we found that less maturation of the superficial white matter was associated with more internalizing behavior problems after injury in females. This suggests that alterations in the trajectory of white matter maturation in females may play a role in the development of new internalizing problems following concussion. The lack of a relationship between deviation from typical white matter maturation and behavior problems in males is consistent with the absence of concussion-induced brain and behavioral effects observed in males. Biological or physiological factors may predispose the female brain to primary and secondary injury effects of concussion and/or reduce the capacity for repair in white matter after a concussion. Potential factors related to sex that impact primary injury include neck strength as greater neck strength in males provides more control over head movements which may decrease the impact of the injury on the brain (Honda, Chang, and Kim 2018). Secondary injury processes after concussion that are known to be sexually dimorphic include microglial activation and neuroinflammation (Kenzie et al. 2017; Armstrong et al. 2016). Additionally, male rodents have been shown to have more myelinated axons, thicker myelin sheaths, larger axon diameters, as well as increased density of oligodendrocytes, and lower rates of oligodendrocyte turnover in white matter compared to females (Dollé et al. 2018; Cerghet et al. 2006; Abi Ghanem et al. 2017) which may contribute to differences in myelin damage and repair in males and females (Cerghet et al. 2009; Mierzwa et al. 2015).

A distinct advantage of examining the impact of concussion on behavior problems and white matter maturation in the ABCD cohort is that it is a large sample of similarly aged children with pre-injury information about behavior and the brain. This allows for the investigation of behavior and brain changes with reduced developmental heterogeneity in the sample. However, there are some important limitations to note. First, concussion history was based on a parent-report measure completed retrospectively with recall from up to two years ago for some participants. Similarly, behavioral problems were based on a parent-report measure and previous studies have shown maternal reports of child behavioral problems can differ with mothers reporting more internalizing behavior problems for female children and more externalizing behavior problems for male children (Najman et al. 2001). Future work should attempt to consider both parent and child reports of behavioral problems. Second, MRI is not specific and the exact microstructural changes influencing increased neurite density are not known. Lastly, the interpretation of our results on brain-behavior associations may be confounded by the small sample of children with concussion that had imaging data available (n = 99). For this reason, follow-up analyses investigating the contribution of injury variables in children with concussion are exploratory.

In conclusion, our findings provide important insights into the contribution of pre-existing behavioral problems and white matter disruption on long-lasting behavioral problems after concussion in children. The unique longitudinal design of the current study that captured pre-concussion measures of behavior and brain allowed for the identification of new behavioral problems and disrupted brain maturation. Sex-specific effects in females were small but consistently significant across analyses. Overall, the results indicated that there is an association between disrupted superficial white matter development and internalizing behaviors in females specifically. This identifies a biological substrate influencing behavioral problems post-concussion. Future therapeutic studies may be able to leverage superficial white matter microstructure as a biomarker in female children with concussion.

