## Supplementary Table 1 for "Disrupted maturation of white matter microstructure after concussion contributes to internalizing behavior problems in female children"

**Supplementary Table 1**. Demographics and injury characteristics of the concussion group with MRI data available

|  | **F (N=35)** | **M (N=64)** | **Total (N=99)** |
| --- | --- | --- | --- |
| **Age at Baseline (in months)** |  |  |  |
| Mean (SD) | 121 (6.67) | 120 (7.13) | 120 (6.94) |
| Median [Min, Max] | 119 [108, 131] | 120 [107, 131] | 120 [107, 131] |
| **Age at Follow-Up (in months)** |  |  |  |
| Mean (SD) | 144 (6.98) | 144 (7.21) | 144 (7.09) |
| Median [Min, Max] | 144 [132, 158] | 146 [129, 156] | 145 [129, 158] |
| **Puberty at Follow-Up** |  |  |  |
| Prepubescence | 12 (34.3%) | 45 (70.3%) | 57 (57.6%) |
| Pubescence | 23 (65.7%) | 19 (29.7%) | 42 (42.4%) |
| **Combined Family Income** |  |  |  |
| <$50K | 4 (11.4%) | 9 (14.1%) | 13 (13.1%) |
| $100K+ | 13 (37.1%) | 35 (54.7%) | 48 (48.5%) |
| $50-99K | 18 (51.4%) | 20 (31.3%) | 38 (38.4%) |
| **Race/Ethnicity** |  |  |  |
| Hispanic | 5 (14.3%) | 10 (15.6%) | 15 (15.2%) |
| Non-Hispanic Black | 3 (8.6%) | 1 (1.6%) | 4 (4.0%) |
| Non-Hispanic White | 25 (71.4%) | 42 (65.6%) | 67 (67.7%) |
| Other/Multi-Racial | 2 (5.7%) | 10 (15.6%) | 12 (12.1%) |
| Asian | 0 (0%) | 1 (1.6%) | 1 (1.0%) |
| **Pre-Injury Internalizing Behaviour Raw Score** |  |  |  |
| Mean (SD) | 7.37 (6.76) | 5.88 (6.12) | 6.40 (6.36) |
| Median [Min, Max] | 6.00 [0, 28.0] | 4.00 [0, 30.0] | 5.00 [0, 30.0] |
| **Pre-Injury Internalizing Behaviour T-Score** |  |  |  |
| Mean (SD) | 52.2 (11.0) | 51.0 (10.9) | 51.4 (10.9) |
| Median [Min, Max] | 52.0 [33.0, 77.0] | 50.0 [34.0, 77.0] | 52.0 [33.0, 77.0] |
| **Pre-Injury Externalizing Behaviour Raw Score** |  |  |  |
| Mean (SD) | 5.54 (5.77) | 5.47 (6.48) | 5.49 (6.21) |
| Median [Min, Max] | 4.00 [0, 24.0] | 3.00 [0, 24.0] | 3.00 [0, 24.0] |
| **Pre-Injury Externalizing Behaviour T-Score** |  |  |  |
| Mean (SD) | 48.8 (10.2) | 47.1 (11.2) | 47.7 (10.9) |
| Median [Min, Max] | 49.0 [34.0, 72.0] | 46.0 [33.0, 71.0] | 47.0 [33.0, 72.0] |
| **Post-Injury Internalizing Raw Behaviour Score** |  |  |  |
| Mean (SD) | 7.06 (6.57) | 5.91 (5.72) | 6.31 (6.03) |
| Median [Min, Max] | 5.00 [0, 26.0] | 3.50 [0, 23.0] | 4.00 [0, 26.0] |
| **Post-Injury Internalizing Behaviour T-Score** |  |  |  |
| Mean (SD) | 51.0 (10.7) | 50.8 (10.4) | 50.9 (10.4) |
| Median [Min, Max] | 50.0 [33.0, 72.0] | 49.0 [34.0, 71.0] | 50.0 [33.0, 72.0] |
| **Post-Injury Externalizing Raw Behaviour Score** |  |  |  |
| Mean (SD) | 4.51 (3.99) | 5.11 (5.36) | 4.90 (4.91) |
| Median [Min, Max] | 3.00 [0, 16.0] | 3.50 [0, 21.0] | 3.00 [0, 21.0] |
| **Post-Injury Externalizing Behaviour T-Score** |  |  |  |
| Mean (SD) | 47.7 (8.20) | 46.9 (9.49) | 47.2 (9.02) |
| Median [Min, Max] | 46.0 [34.0, 64.0] | 47.0 [33.0, 69.0] | 46.0 [33.0, 69.0] |
